# Accurate recognition of colorectal cancer with semi-supervised deep learning on pathological images

**DOI:** 10.1101/2020.07.13.201582

**Authors:** Gang Yu, Ting Xie, Chao Xu, Xing-Hua Shi, Chong Wu, Kai Sun, Run-Qi Meng, Xiang-He Meng, Kuan-Song Wang, Hong-Mei Xiao, Hong-Wen Deng

## Abstract

**Background:** Machine-assisted recognition of colorectal cancer (CRC) has been mainly focused on supervised deep learning that suffers from a significant bottleneck of requiring massive labeled data. We hypothesize that semi-supervised deep learning leveraging a small amount of labeled data with abundant available unlabeled data can provide a powerful alternative strategy.

**Method:** We proposed a semi-supervised model based on the mean teacher architecture that provides pathological predictions at both patch- and patient-levels. We demonstrated the general utility of the model utilizing 13,111 CRC whole slide images from 8,803 subjects gathered from 13 independent centers. We compared our proposed method with the prevailing supervised learning and six pathologists. Two extended evaluations on 15,000 lung and 294,912 lymph node images were also performed to confirm the generality of utility of semi-supervised learning for different cancers.

**Results:** With a small amount of labeled training patches (∼3,150 labeled, ∼40,950 unlabeled or ∼6,300 labeled, ∼37,800 unlabeled), the semi-supervised learning (SSL) performed significantly better than the supervised learning (SL, which only used the labeled data) (area under the curve, AUC: 0.90 ± 0.06 vs. 0.84 ± 0.07, *P* value = 0.02 or AUC: 0.98 ± 0.01 vs. 0.92 ± 0.04, *P* value = 0.0004). Moreover, we found no significant difference between SL using massive ∼44,100 labeled patches and SSL (∼6,300 labeled, ∼37,800 unlabeled) at patch-level diagnoses (AUC:0.98 ± 0.01 vs. 0.987 ± 0.01, *P* value = 0.134) and patient-level diagnoses (average AUC: 0.974 vs. 0.980, *P* value = 0.117). SSL was close to human pathologists in diagnosis performance (average AUC: 0.972 vs. 0.969). This extended evaluation on lung and lymph node also confirmed when a small amount of labeled data were used, SSL was better than SL, and achieved similar performance as that of SL with massive labeling.

**Conclusions:** We reported that SSL can achieve excellent performance through a multi-center study. Because SSL dramatically reduces the need and cost of pathological image annotation, it has great potential to effectively build pathological artificial intelligence (AI) platforms in practice.

## Introduction

Colorectal cancer (CRC) is the second most common cause of cancer death in Europe and America [1-2]. Pathological diagnosis is one of the most authoritative methods for diagnosing CRC [3-4], which requires a pathologist to visually examine digital full-scale whole slide images (WSI). The challenges stem from the complexity of WSI including large image sizes (> 10,000 × 10,000 pixels), complex shapes, textures, and histological changes in nuclear staining [4]. Furthermore, there is a shortage of pathologists worldwide in stark contrast with the rapid accumulation of WSI data, and the daily workload of pathologists is intensive which could lead to unintended misdiagnose due to fatigue [5]. Hence, it is crucial to develop diagnosing strategies that are effective yet of low cost by leveraging recent AI development.

Deep learning provides an exciting opportunity to support and accelerate pathological analysis [6,25], including lung [7-8], breast [9], lymph node [10], and skin cancers [11-12]. Progress has been made in applying deep learning to CRC including classification [13], tumor cell detection [14-15], and outcome prediction [16-18]. We have developed a recognition system for CRC using SL (supervised learning), which achieved one of the highest diagnosis accuracies in cancer diagnosis with AI [19]. However, our earlier method was built upon learning from 62,919 labeled patches from 842 subjects, which were carefully selected and extensively labeled by pathologists.

While SL with massive labeled data can achieve high diagnostic accuracy, the reality is that we often have only a small amount of labeled data and a much larger amount of unlabeled data in medical domains. Although unsupervised learning does not require any labeled data, its performance is still limited currently [20,21]. There are some other approaches for learning on the small amount of labeled data. For example, in transfer learning, the network is firstly trained in a big dataset of source domain, and then trained in labeled medical images. However, the number of labeled images needed is still quite large [22, 23]. The generative adversarial networks (GAN) can generate large amount of data by learning the style from a limited dataset [24, 25]. These approaches may improve accuracy, but they only used limited labeled datasets, and large amounts of unlabeled data do appear in medical domains and clinical settings. Moreover, it would be difficult for GAN to simulate all possible features of the disease based on limited samples.

The SSL (semi-supervised learning), a method that leverages both labeled and unlabeled data is supposed to provide a low-cost alternative in terms of the requirement of the laborious and sometimes impractical sample labeling [26, 27]. Although SSL can improve accuracy of natural images, its performance on medical images is unclear. Recently, some studies were proposed to determine whether SSL based on a small amount of labeled data and a large amount of unlabeled data can improve medical image analysis [28-30], such as object detection [31], data augmentation [32], image segmentation [33,34]. However, only very limited few studies have investigated if SSL can be applied to achieve satisfactory accuracy in pathological images [35], where on a small data set of 115 WSIs, an SSL method of CRC recognition can achieve best accuracy of 0.938 only at 7,180 patches of 50 WSIs from one data center, suggesting the potential of SSL for diagnosis on *patch-level*.

However, to the best of our knowledge, the CRC recognition system of SSL has not been extensively validated on *patient level* dataset from multiple centers to assess the general utility of SSL. How to translate the patch-level prediction to WSI and patient level diagnosis is not trivial. Because we and other groups have not been able to develop perfect patch-level models, the errors at patch-level may be easily magnified on WSI level diagnosis. For example, even though the imperfect patch-level model may yield reasonable prediction on positive (cancerous) WSIs, it also may yield high false positive errors on the negative (non-caner) WSIs, because the false positive errors at patch level will accumulate due to the testing of multiple patches in WSI. However, the patient-level diagnosis is required in clinical applications of any AI system for cancer diagnosis.

To fill this gap, we used 13,111 WSIs collected from 8,803 subjects from 13 independent centers to develop a semi-supervised model. We evaluated SSL by comparing its performance with that of prevailing SL and also with that of professional pathologists. To confirm that SSL can achieve excellent performance on pathological images and further demonstrate our main point that a reliable medical AI can be built with a small amount of labeled data plus other available unlabeled daya, we evaluated it on two other types of cancer (lung cancer and lymphoma). The main contributions of this study are summarized as follows:

1. We evaluated different CRC recognition methods based on SSL and SL at the patch-level and patient-level respectively. This large-scale evaluation showed that accurate CRC recognition is feasible with a high degree of reliability even when the amount of labeled data is limited.
2. We found that when ∼6,300 labeled patches (assuming a large number of unlabeled patches (e.g., ∼37,800) available, which was often the case in practice) were used for SSL, there was no significant difference between SSL and SL (developed based on ∼44,100 labeled patches) and pathologists. This finding holds for CRC recognition at both the patch level and patient level.
3. The extended experiment of lung cancer and lymphoma further confirmed the conclusion that when a small amount of labeled data plus a large amount of unlabeled data were used, SSL may perform similarly or even better than SL. Our study thus indicated that SSL will dramatically reduce the amount of labeled data required in practice, to greatly facilitate the development and application of AI in medical sciences.

## Methods

We trained and tested our method utilizing CRC datasets from multiple centers (Table 1), which were divided into four big datasets for different aims (Dataset-PATT, Dataset-PAT, Dataset-PT, Dataset-HAC, Supplementary Table 1). Briefly, we divided each WSI into thousands of patches. At the patch level, a patch-level model was used for the prediction of cancer or non-cancer patch. For simplicity, we used SSL, SL to represent semi-supervised and supervised learning methods, and a numerical number to represent labels used in training set, which accounts for the percentage of 62,919 patches in Dataset-PATT (Table 2). Five versions of patch-level model, Model-n%-SSL or Model-n%-SL were trained based on different training sets and learning method (Table 3, Figure 1 (a)) and then tested (Figure 1 (b)). For SL, Inception V3 [36] was used for patch-level model. For SSL, we applied an SSL strategy called mean teacher [26], where two Inception V3 networks, teacher network and student network were trained.

**Table 1.**
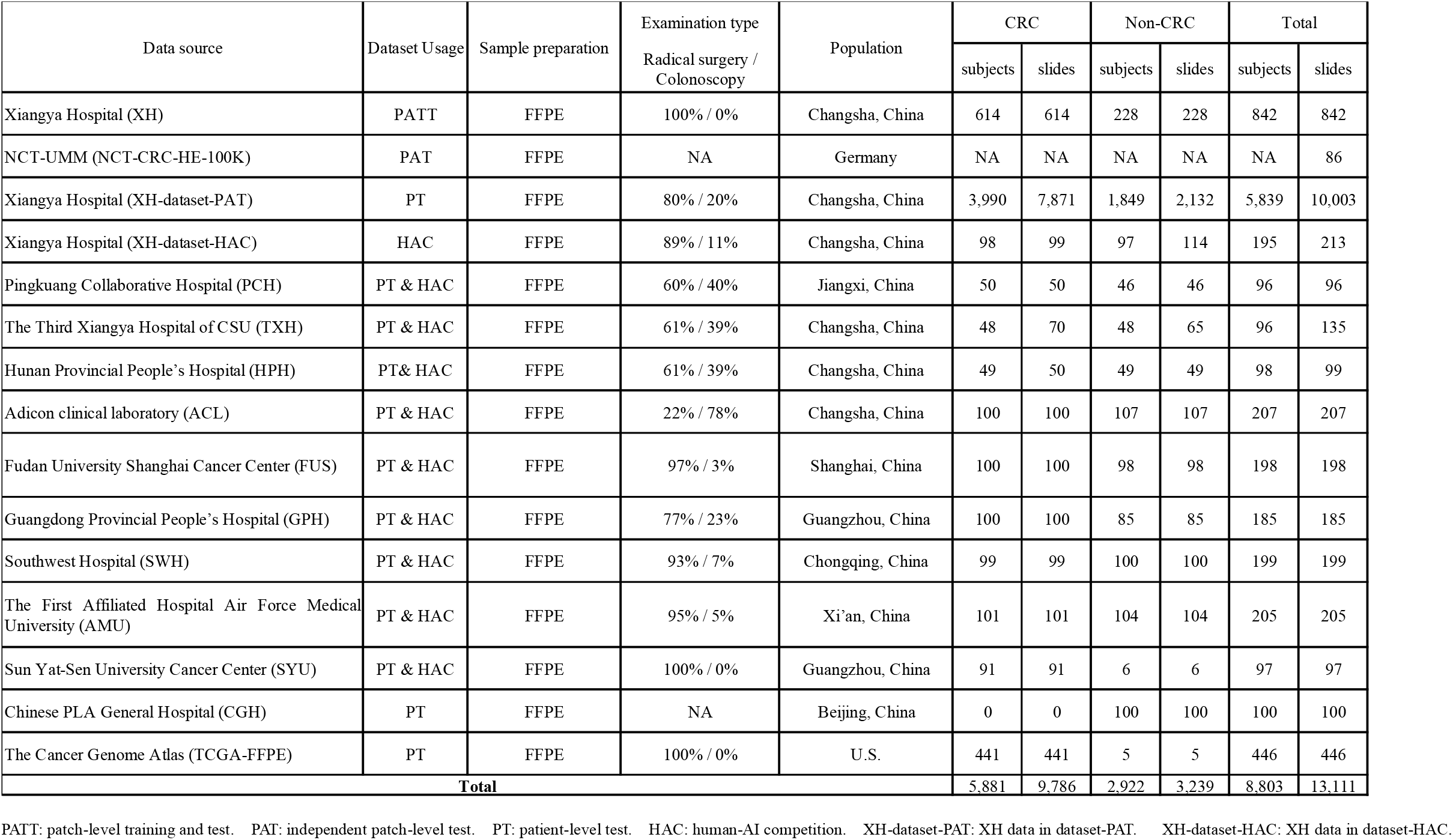
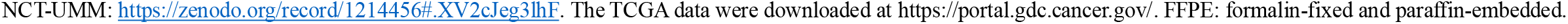
Datasets used from multi-center data sources.

| Data source | Dataset Usage | Sample preparation | Examination type<br>Radical surgery /<br>Colonoscopy | Population | CRC |  | Non-CRC |  | Total |  |
| --- | --- | --- | --- | --- | --- | --- | --- | --- | --- | --- |
|  |  |  |  |  | subjects | slides | subjects | slides | subjects | slides |
| Xiangya Hospital (XH) | PATT | FFPE | 100% / 0% | Changsha, China | 614 | 614 | 228 | 228 | 842 | 842 |
| NCT-UMM (NCT-CRC-HE-100K) | PAT | FFPE | NA | Germany | NA | NA | NA | NA | NA | 86 |
| Xiangya Hospital (XH-dataset-PAT) | PT | FFPE | 80% / 20% | Changsha, China | 3,990 | 7,871 | 1,849 | 2,132 | 5,839 | 10,003 |
| Xiangya Hospital (XH-dataset-HAC) | HAC | FFPE | 89% / 11% | Changsha, China | 98 | 99 | 97 | 114 | 195 | 213 |
| Pingkuang Collaborative Hospital (PCH) | PT & HAC | FFPE | 60% / 40% | Jiangxi, China | 50 | 50 | 46 | 46 | 96 | 96 |
| The Third Xiangya Hospital of CSU (TXH) | PT & HAC | FFPE | 61% / 39% | Changsha, China | 48 | 70 | 48 | 65 | 96 | 135 |
| Hunan Provincial People's Hospital (HPH) | PT & HAC | FFPE | 61% / 39% | Changsha, China | 49 | 50 | 49 | 49 | 98 | 99 |
| Adicon clinical laboratory (ACL) | PT & HAC | FFPE | 22% / 78% | Changsha, China | 100 | 100 | 107 | 107 | 207 | 207 |
| Fudan University Shanghai Cancer Center (FUS) | PT & HAC | FFPE | 97% / 3% | Shanghai, China | 100 | 100 | 98 | 98 | 198 | 198 |
| Guangdong Provincial People's Hospital (GPH) | PT & HAC | FFPE | 77% / 23% | Guangzhou, China | 100 | 100 | 85 | 85 | 185 | 185 |
| Southwest Hospital (SWH) | PT & HAC | FFPE | 93% / 7% | Chongqing, China | 99 | 99 | 100 | 100 | 199 | 199 |
| The First Affiliated Hospital Air Force Medical University (AMU) | PT & HAC | FFPE | 95% / 5% | Xi'an, China | 101 | 101 | 104 | 104 | 205 | 205 |
| Sun Yat-Sen University Cancer Center (SYU) | PT & HAC | FFPE | 100% / 0% | Guangzhou, China | 91 | 91 | 6 | 6 | 97 | 97 |
| Chinese PLA General Hospital (CGH) | PT | FFPE | NA | Beijing, China | 0 | 0 | 100 | 100 | 100 | 100 |
| The Cancer Genome Atlas (TCGA-FFPE) | PT | FFPE | 100% / 0% | U.S. | 441 | 441 | 5 | 5 | 446 | 446 |
| **Total** |  |  |  |  | **5,881** | **9,786** | **2,922** | **3,239** | **8,803** | **13,111** |

PATT: patch-level training and test. PAT: independent patch-level test. PT: patient-level test. HAC: human-AI competition. XH-dataset-PAT: XH data in dataset-PAT. XH-dataset-HAC: XH data in dataset-HAC.
NCT-UMM: https://zenodo.org/record/1214456#.XV2cJeg3lhF. The TCGA data were downloaded at https://portal.gdc.cancer.gov/. FFPE: formalin-fixed and paraffin-embedded.

**Table 2.** Dataset-PATT and Dataset-PAT

| Dataset | Cancer |  |  | Non-cancer |  |  | Total |  |  |
| --- | --- | --- | --- | --- | --- | --- | --- | --- | --- |
|  | subjects | slides | patches | subjects | slides | patches | subjects | slides | patches |
| **Dataset-PATT** | 614 | 614 | 30,056 | 228 | 228 | 32,863 | 842 | 842 | 62,919 |
| **Dataset-PAT<sup>a</sup>** | NA | NA | 14,317 | NA | NA | 85,683 | NA | 86 | 100,000 |
| **Total** | >614 | >614 | 44,373 | >228 | >228 | 118,546 | >842 | 928 | 162,919 |

a: No information (NA) on the number of cancer or non-cancer subjects/slides was provided.

**Table 3.** Training and testing set for CRC patch-level models

| Model | Class | Dataset-PATT |  |  | Dataset-PAT |
| --- | --- | --- | --- | --- | --- |
|  |  | Training set |  | Testing set |  |
|  |  | Labeled | Unlabeled<sup>f</sup> |  |  |
| Model-5%-SSL | Cancer | 1,645 | 21,390 | 9,828 | 14,317 |
|  | Non-cancer | 1,505 | 19,560 | 8,991 | 85,683 |
|  | Total | 3,150/5%<sup>a</sup> | 40,950/65%<sup>b</sup> | 18,819/30%<sup>c</sup> | 100,000 |
| Model-10%-SSL | Cancer | 3,290 | 19,745 | 9,828 | 14,317 |
|  | Non-cancer | 3,010 | 18,055 | 8,991 | 85,683 |
|  | Total | 6,300/10% | 37,800/60%<sup>d</sup> | 18,819/30% | 100,000 |
| Model-5%-SL | Cancer | 1,645 | - | 9,828 | 14,317 |
|  | Non-cancer | 1,505 | - | 8,991 | 85,683 |
|  | Total | 3,150/5% | - | 18,819/30% | 100,000 |
| Model-10%-SL | Cancer | 3,290 | - | 9,828 | 14,317 |
|  | Non-cancer | 3,010 | - | 8,991 | 85,683 |
|  | Total | 6,300/10% | - | 18,819/30% | 100,000 |
| Model-70%-SL | Cancer | 23,035 | - | 9,828 | 14,317 |
|  | Non-cancer | 21,065 | - | 8,991 | 85,683 |
|  | Total | 44,100/70%<sup>e</sup> | - | 18,819/30% | 100,000 |

a-e: Because the number of patches from each WSI is not the same, the number of patches estimated based on the proportion of extraction is approximate.
f: the labels of the patches are ignored.

**Figure 1.**
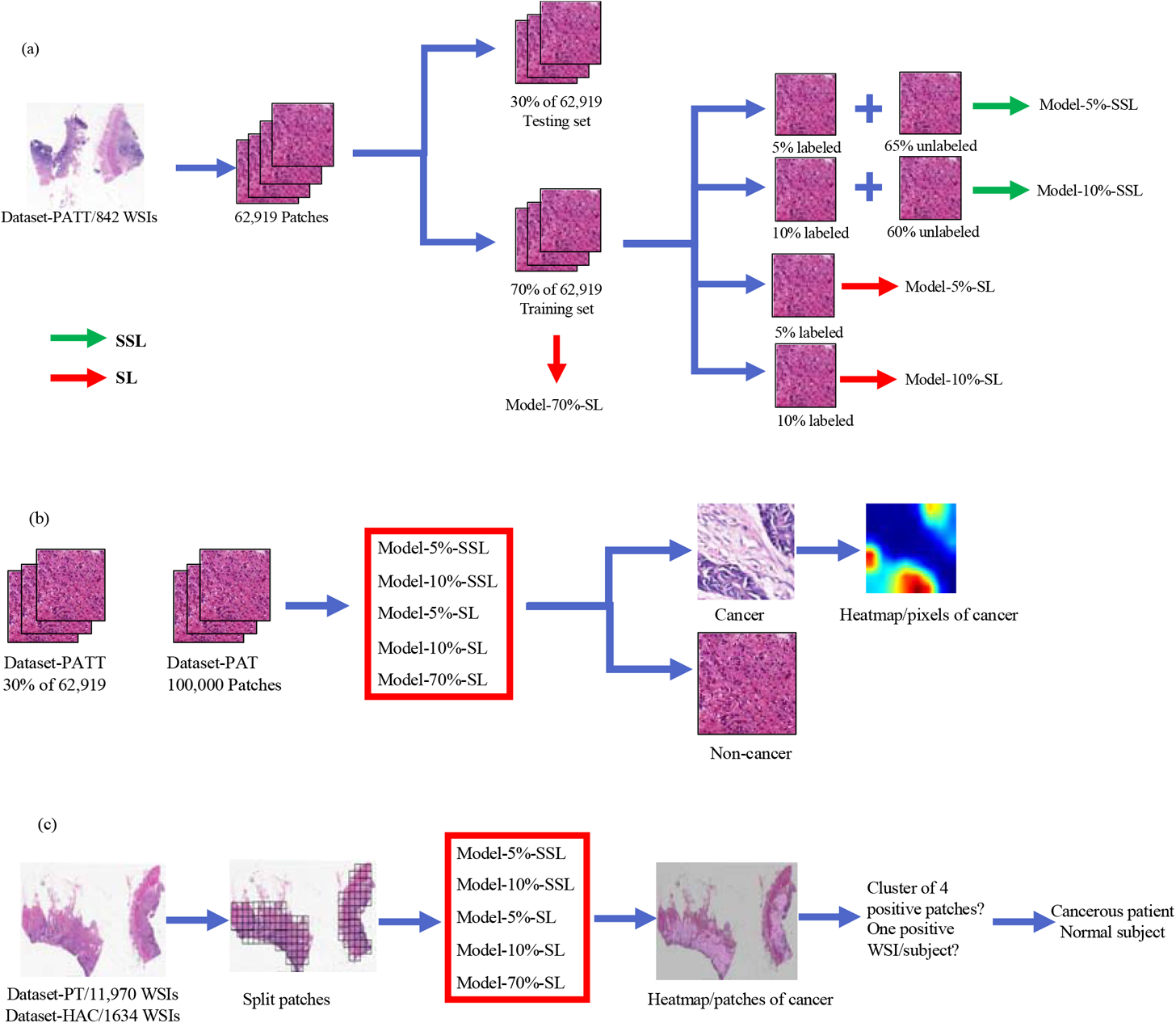
The flow chart of the CRC study. (a) SSL and SL are performed on different labeled and unlabeled patches from Dataset-PATT. (b) The patch-level test on 30% of Dataset-PATT and whole dataset of Dataset-PAT. (c) The patient-level test of Dataset-PT and Dataset-HAC. If there is a cluster of four positive patches on WSI, the WSI is positive. A subject with a positive WSI is cancerous.

We aimed to evaluate two hypotheses at the patch level. First, we hypothesized that SSL is better than SL when only a small number (∼thousands) of labeled patches available for both SL and SSL and a large number (∼ten of thousands) of unlabeled patches are also available for SSL. Second, we further hypothesized that there is no significant difference between SSL using a few thousands of labeled patches (plus a larger number of unlabeled patches) and SL using massive labeled patches (tens of thousands).

At the WSI and patient levels, we applied a cluster-based and positive sensitivity strategy to achieve CRC diagnosis for patients as we did recently [19] (Figure 1). To save space, the details of the CRC diagnosis methodology were relegated to Supplementary A, B. The source code and data can be found: www.github.com/csu-bme/pathology_SSL.

## Results

### SSL vs SL CRC recognition at patch level

The 62,919 patches in Dataset-PATT were used for patch-level training and testing (Table 3, Figure 1). The 30% of 62,919 (∼18,819) were used for the testing, and the remaining 70% of the patches (44,100) were used for the training. Model-5%-SSL and Model-10%-SSL were trained on 5% (∼3,150) and 10% (∼6,300) labeled patches, respectively, where the remaining patches 65% (∼40,950) and 10% (∼37,800) were used, but their labels were ignored (as unlabeled patches). Model-5%-SL and Model-10%-SL were trained on the same labeled patches (5%, ∼3,150 and 10%, ∼6,300) only with Model-5%-SSL and Model-10%-SSL respectively, but the remained patches (40,950, 37,800) were not used. Model-70%-SL was trained on the ∼44,100 labeled training patches.

The area under the curve (AUC) distribution on Dataset-PATT and Dataset-PAT were shown in Figure 2. Model-5%-SSL was superior to Model-5%-SL (AUC of both Dataset-PATT and Dataset-PAT, 0.75 confidence interval: 0.90±0.06 vs.0.84±0.07, P value=0.02, Wilcoxon signed rank test). Model-10%-SSL was also significantly better than Model-10%-SL (AUC: 0.98±0.01vs. 0.92±0.04, P value=0.0004). These results indicated that when approximately 3,150 or 6,300 patches were labeled, the SSL method was always better than SL.

**Figure 2.**
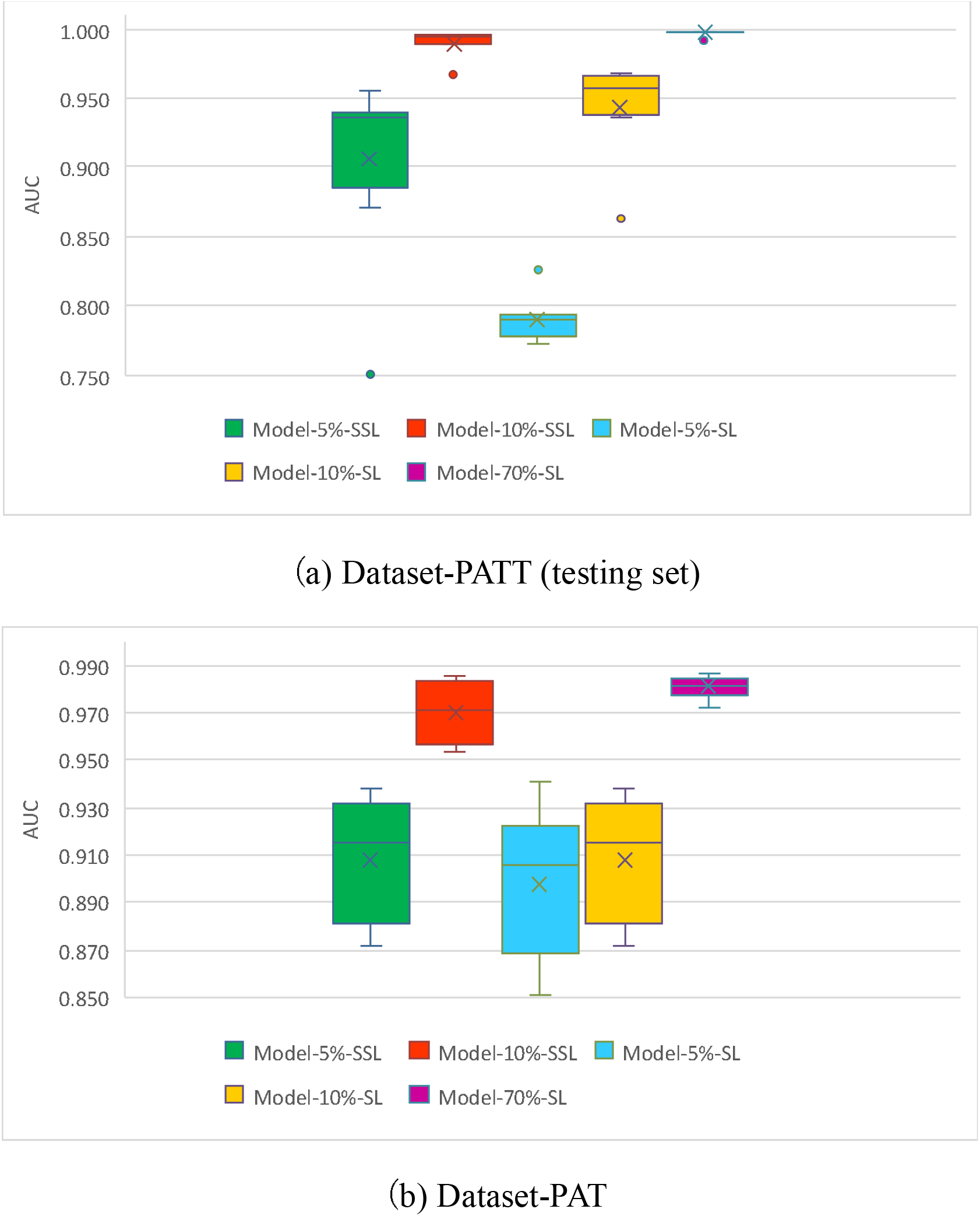
AUC distribution of five models at patch level on Dataset-PATT and Dataset-PAT. The horizontal bar in the box indicates the median, while the cross indicates the mean.

The performance of Model-10%-SSL had no significant difference with that of Model-70%-SL (AUC: 0.98±0.01 vs. 0.987±0.01, P value=0.134). This observation indicated that there was no significant difference between the SSL method (6,300 labeled, 37,800 unlabeled) and the SL (44,100 labeled). Visual inspection (Supplementary Figure 2) confirmed that that Model-10%-SL could not really find the locations of cancer in the patches, while the locations of cancer by Model-10%-SSL and Model-70%-SL were highly matched.

### Patient-level CRC recognition

To test whether the above conclusion at patch-level still holds at patient-level, we evaluated three of five models using Dataset-PT (Figure 3 and Supplementary C). The patient-level diagnosis is based on the recognition of every patch provided by patch-level models, and then cluster-based WSI inference and positive sensitivity for patient inference (Figure 1(c)).

**Figure 3.**
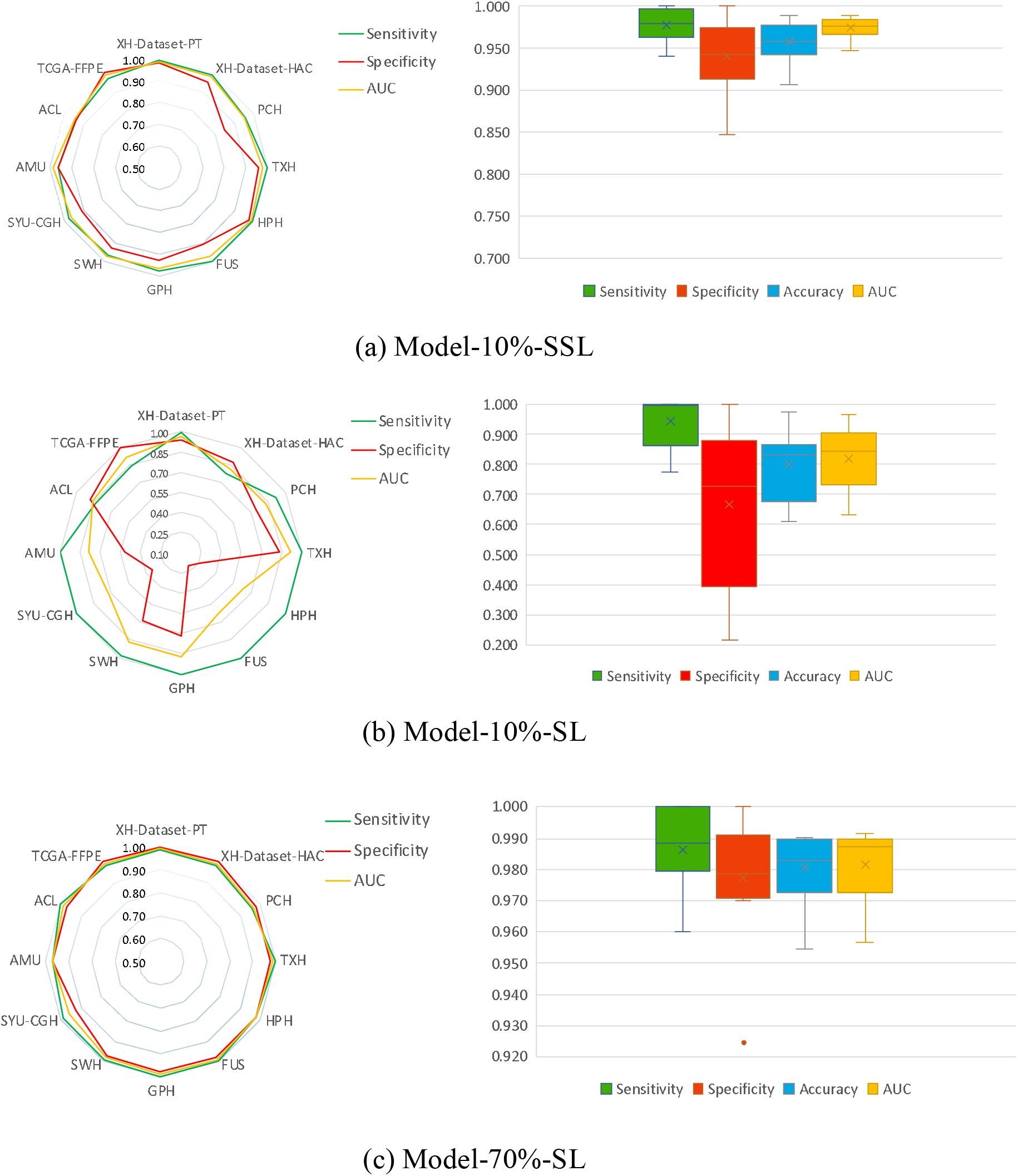
Patient-level comparison on twelve independent datasets. Left: Radar maps illustrating the sensitivity, specificity, and AUC. Right: Boxplots showing the distribution of sensitivity, specificity, accuracy, and AUC in these datasets.

Model-10%-SSL had a significant improvement over Model-10%-SL (Average AUC: 0.974 vs. 0.819, *P* value = 0.0022) on patient-level prediction in the multi-centers scenario. The average AUC of Model-10%-SSL was slightly lower than, but comparable to, that of Model-70%-SL (average AUC: 0.974 vs. 0.980, *P* value = 0.117). Among the 7 datasets (XH-dataset-PT, XH-dataset-HAC, PCH, TXH, FUS, SWH, TCGA, 11,290 WSIs), the AUC difference of Model-10%-SSL and Model-70%-SL was smaller than 1.6%. In particular, on the largest dataset, XH-dataset-PT (10,003 WSIs), the AUCs of Model-10%-SSL and Model-70%-SL were close with 0.984 vs. 0.992. On the datasets of HPH, SYU, CGH and AMU (501 WSIs), the AUCs of Model-10%-SSL were even higher than that of Model-70%-SL.

In the data from GPH, and ACL (392 WSIs), the performance of Model-10%-SSL was lower than that of Model-70%-SL (AUC DIFF>0.222). It is worth noting that Model-10%-SSL generally achieved good sensitivity, which proved practically useful for the diagnosis of CRC. Visual inspection in Supplementary Figure 3 showed the cancer patches identified by Model-10%-SSL and Model-70%-SL were the true cancer locations on WSIs.

### Human-AI CRC competition

We recruited six pathologists with 1-18 years of independent experience (Supplementary Table 4). They independently reviewed 1,634 WSIs from 10 data centers (Dataset-HAC, Figure 4) with no time limit and diagnosed the cancer solely based on WSIs (i.e., no other clinical data were used). We ranked pathologists, Model-10%-SSL and Model-70%-SL. The average AUC of model-10%-SSL was 0.972, ranked at the 5th, which was close to the average AUC of pathologists (0.969). The sensitivity of Model-10%-SSL was 0.977, showing an excellent detection ability of cancer (Supplementary Table 5).

**Figure 4.**
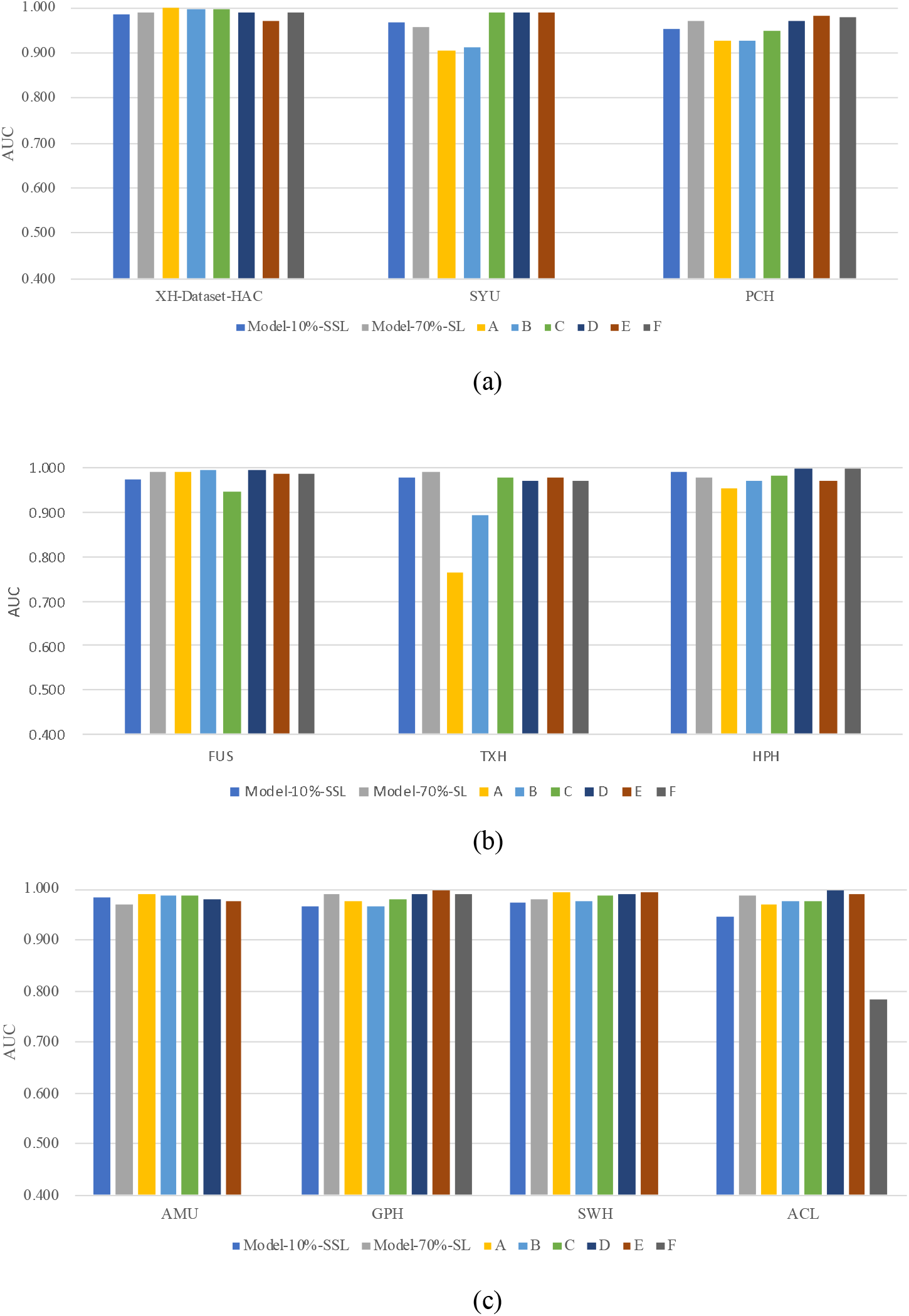
AUC comparison of in the Human-AI contest using Dataset-HAC, which consists XH-dataset-HAC, PCH, TXH, HPH, ACL, FUS, GPH, SWH, AMU and SYU. Colored lines indicate the AUCs achieved by THE two models and six pathologists (A-F). The F pathologist didn’t attend the competition of SYU, AMU and SWH dataset.

### Extended SSL vs SL experiment of lung and lymph node cancer

In order to demonstrate the utility of SSL on other pathological images, the experiment of SSL and SL on lung and lymph node were performed as detailed in Supplementary D for Methodology. 15,000 lung images of three classes: adenocarcinoma, squamous cell carcinoma, and benign tissue were obtained from LC25000 dataset (Lung) [38], and the 294,912 lymph node images including tumor and benign tissue were obtained from PatchCamelyon (Pcam) [39]. Similarly, SSL was trained on a small number of labeled images and a large number of unlabeled images (for which labels are known but ignored during training), and compared with SL (Supplementary Tables 6 and 7). The base model of SL is Inception V3, and mean teacher method is also used for SSL, repeatedly trained 8 times as in the main CRC study to assess various performance statistics.

Because the number of classes in lung images is three, the accuracy is used for the evaluation. Lung-5%-SSL (5% labeled and 75% unlabeled) and Lung-20%-SSL (20% labeled, 60% unlabeled) were better than Lung-5%-SL (5% labeled) and Lung-20%-SL (20% labeled) (Accuracy:0.960±0.007 vs 0.918±0.026, *P* value=0.012; 0.989±0.003 vs 0.961±0.025, *P* value=0.011, Figure 5) respectively. There was no difference between Lung-20%-SSL and Lung-80%-SL (80% labeled) (Accuracy: 0.989±0.003 vs 0.993±0.002, *P* value=0.093). Pcam-1%-SSL (1% labeled, 99% unlabeled) and Pcam-5%-SSL (5% labeled, 95% unlabeled) are better than Pcam-1%-SL (1% labeled) and Pcam-5%-SL (5% labeled) (AUC: 0.947±0.01 vs 0.912±0.01, *P* value=0.012; 0.960±0.002 vs 0.943±0.01, *P* value=0.0001, Figure 6) respectively. Pcam-5%-SSL can be compared to Pcam-100%-SL (100% labeled) (AUC: 0.960±0.002 vs 0.961±0.005, *P* value=0.94). This extended experiment confirmed the conclusion that when a small number of labeled pathological images was available together with a large number of unlabeled image data, SSL can be compared to SL with massive labels.

**Figure 5.**
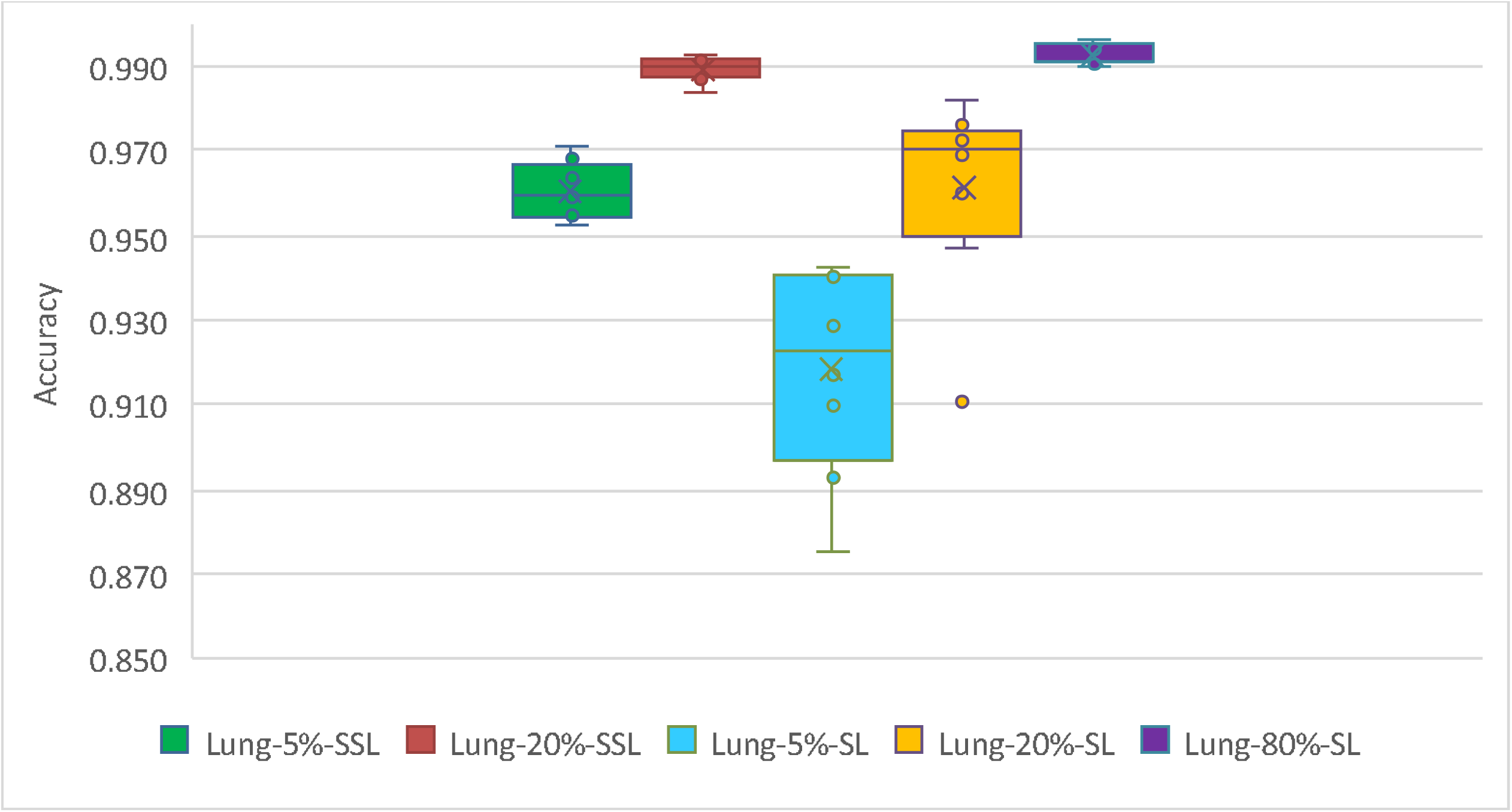
Accuracy distribution of five models on lung.

**Figure 6.**
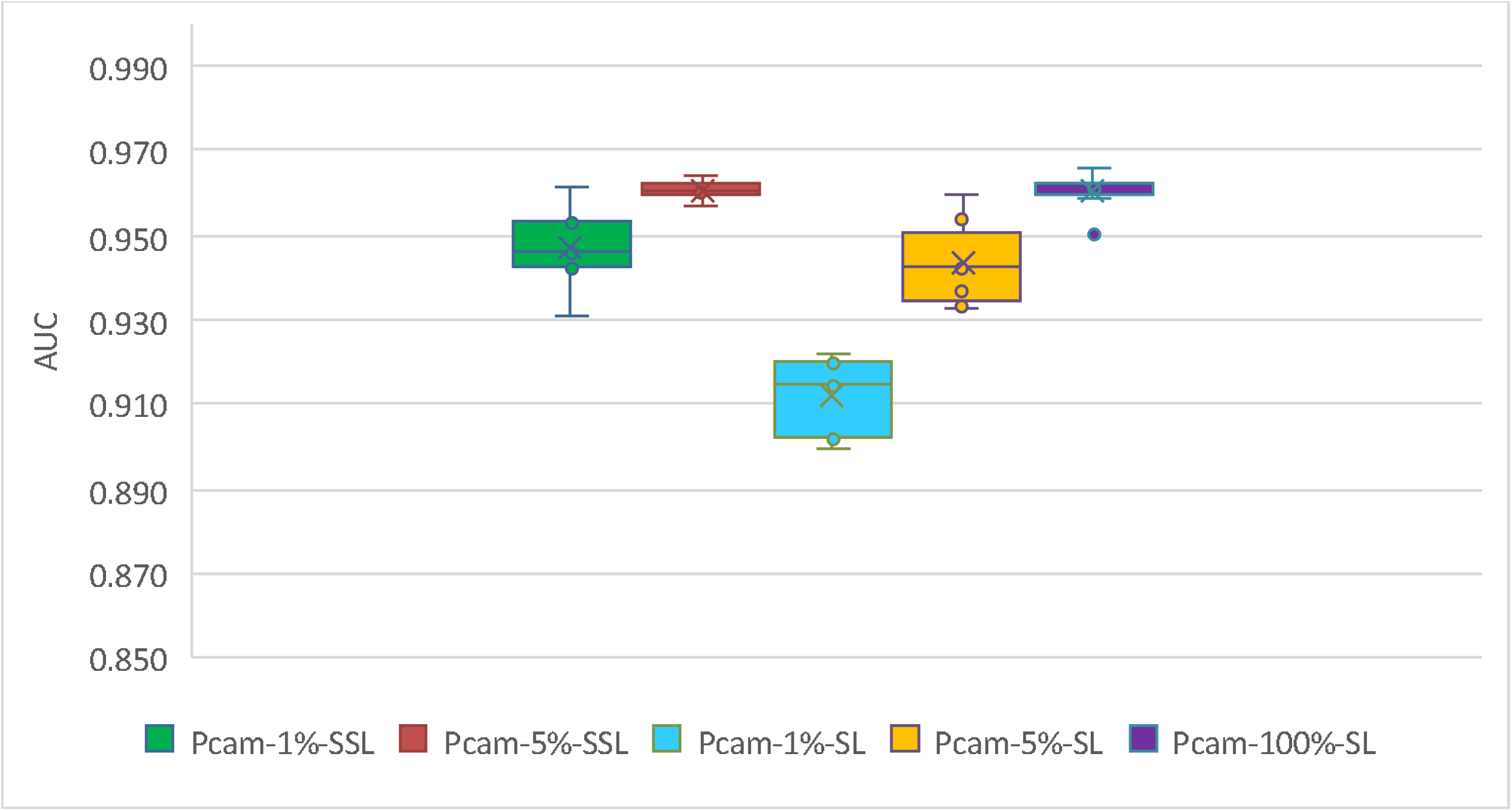
AUC distribution of five models on lymph node.

### Comparison with related research

We compared our methods with seven existing CRC detection methods [35,40-45], and five other cancers (lung, ductal carcinoma, breast, prostate, basal cell carcinoma) detection methods [46-50] (Supplementary Table 10). The 6 of 7 CRC detection methods had an AUC ranging from 0.904 to 0.99 based on SL. Besides, Shaw et al [35] used cancer and normal patches in 86 CRC WSIs to develop a SSL detection method, and used the test set of 7,180 patches in 50 WSIs with colorectal adenocarcinoma, all from one data center, with the best accuracy of 0.938 confirming the potential of SSL on patch-level. In this study, we showed the advantages of the SSL method with 162,919 patches and 13,111 WSIs at both patch and patient levels from multiple independent centers, attesting to the robustness and general utility of the SSL model we developed, where the Model-10%-SSL is comparable to the recent SL models [19]. Besides, Lung-20%-SSL is also comparable to the SL of Coudray et al for lung cancer detection [46].

## Discussion

Accurately diagnosing CRC requires years of training, leading to a global shortage of pathologists [2]. Almost all existing computer-assisted diagnosis models currently relies on massive labeled data with SL, but manual labeling is usually time-consuming and costly. This leads to an increasing interest in building an accurate diagnosis system with far less labeled data.

In this study, we applied SSL to CRC diagnosis, and evaluated its performance using an extensive collection of WSIs across 13 medical centers. On this large data set, we conducted a range of comparison of CRC recognition performance among SSL, SL and six human pathologists, at both patch level and patient level.

We demonstrated that SSL outperformed SL at patch-level recognition when only a small amount of labeled and large amounts of unlabeled data were available. In our previous study [19], we used 62,919 labeled patches from 842 WSIs, which achieved accurate patch-level recognition. When SSL was used as demonstrated in this study, only about a tenth (6,300) of those many labeled patches plus 37,800 unlabeled patches were used to achieve similar AUC to [19] (i.e. Model-70%-SL).

We also conducted extensive testing of three models for patient level prediction on 12 centers (Dataset-PT). Just like the patch level, at the patient level, the SSL outperformed the SL when a small number of labeled patches was available, and close to SL when using a large number of labeled patches. The AUC of Model-10%-SL at XH-Dataset-PT was 0.964, perhaps because both the testing data and training data were from XH.

However, using the data from 12 centers, the average AUC of Model-10%-SL was dramatically reduced to 0.819 from 0.964. This result showed that when training data and testing data were not the same source, the generalization performance of Model-10%-SL was significantly reduced. Moreover, many cancerous patches predicted by Model-10%-SL was deviated from true cancer locations in a WSI (Supplementary Figure 3).

When a large number of unlabeled patches was added for SSL, the generalization performance across centers can be maintained, where there was no significant difference when comparing with Model-70%-SL using massive labeled patches. These results showed that when labeled patches were seriously insufficient, using unlabeled data can greatly improve the generalization ability across different data sets. The patient-level results indicated that with SSL, we may not need as much labeled data as in SL. Since it is well known that unlabeled medical data are relatively easy to obtain, it is of great importance and with an urgent need to develop SSL methods.

We compared the diagnosis of six pathologists from SSL. We found that SSL reached an average AUC of pathologists, which was approximately equivalent to a pathologist with five years of clinical experience. The Human-AI competition in this regard thus showed that it was feasible to build an expert-level method for clinical practice based on SSL. Based on the extended experiments of cancers of lung and lymph node, we further confirmed the conclusion on CRC that when a small amount of labeled data was used, SSL plus a large amount of unlabeled data performed better than SL (with the same number of labeled images). SSL performance can be compared to SL with massive labeling, which confirms the conclusion that SSL may reduce need for the amount of annotation data on pathological images.

In practice, the exact amount of the data that needs to be labeled is generally unknown. Nonetheless, as shown in our experiments, it is an alternative low-cost approach to conduct SSL. Hence, it is an effective strategy to wisely utilize all data so that a small amount of data is first labeled to build a baseline model based on SSL. If the results are not satisfactory for this baseline model, the amount of labeled data should be increased. This strategy is feasible since as expected, SSL requires a much smaller amount of labeled data to achieve the same performance compared with SL.

Although studies have shown that SSL achieved good results in tasks like natural image processing, SSL has not been widely evaluated for analyzing pathological images. Our work confirmed that unlabeled data could improve the accuracy on insufficient labeled pathological images. SSL may have excellent potentials to overcome the bottleneck of insufficient labeled data as in many medical domains.

## Conclusion

Currently, patient-level computer-assisted pathological diagnosis is solely based on SL, which requires a large amount of labeled data to achieve good performance. In this study, we applied SSL and extensively evaluated its performance on multi-center datasets. We demonstrated that SSL with a small amount of labeled data of three cancers achieved comparable prediction accuracy as that of SL with massive labeled data and that of experienced pathologists. This study thus supported potential applications of SSL to develop medical AI systems.

## Supporting information

supple-Accurate recognition of colorectal cancer

## Acknowledgement

G.Y. was supported by the Emergency Management Science and Technology Project of Hunan Province (#2020YJ004, #2021-QYC-10050-26366). K.S.W was supported by the National Natural Science Foundation of China (#81673491), Natural Science Foundation of Hunan Province (#2015JJ2150). H.M.X was supported by the National Key Research and Development Plan of China (2017YFC1001103, 2016YFC1201805), National Natural Science Foundation of China (#81471453), and Jiangwang Educational Endowment. H.W.D. were partially supported by the National Institutes of Health (R01AR059781, P20GM109036, R01MH107354, R01MH104680, R01GM109068, R01AR069055, U19AG055373, R01DK115679), the Edward G. Schlieder Endowment and the Drs. W.C.Tsai and P.T.Kung Professorship in Biostatistics from Tulane University.

## Data Availability

The minimum datasets can be found in the link: https://doi.org/10.6084/m9.figshare.15072546.v1. More data can be found from the readme file in the link: www.github.com/csu-bme/pathology_SSL.

## Code Availability

The source code can be found: www.github.com/csu-bme/pathology_SSL.

