## Supplementary material for "Accurate recognition of colorectal cancer with semi-supervised deep learning on pathological images": supple-Accurate recognition of colorectal cancer

**Supplementary Files**

A. **CRC Semi-supervised Learning Methodology**

**Datasets**

Our CRC dataset was composed of 13,111 WSIs collected from 13 sources, including 10 hospitals, a professional clinical laboratory (ACL), two public databases (Table 1). The CRC WSIs were then divided into four datasets for different aims (Dataset-PATT, Dataset-PAT, Dataset-PT, Dataset-HAC, Supplementary Table 1). All WSIs were made from formalin-fixed and paraffin-embedded (FFPE) method.

Dataset-PATT was used for patch-level training and testing, Dataset-PAT for independent patch-level test. All the images from other hospitals as well as ACL (Dataset-PT) were used for patient-level testing. Dataset-HAC were used for human-AI competition.

Dataset-PATT included 62,919 patches (cancer 30,056, non-cancer 32,863) from 842 subjects (cancer 614, non-cancer 228, Table 2) from Xiangya Hospital (XH). The Dataset-PAT (NCT-CRC-HE-100K) from NCT biobank and the UMM pathology archive (NCT-UMM, National Center for Tumor diseases, University Medical Center Mannheim, Heidelberg University, Germany) was used for further patch-level validation, where there were 100,000 patches from 86 slides of CRC tissue. All the patches can be downloaded at <https://zenodo.org/record/1214456#.XV2cJeg3lhF>, whose labels were from the NCT-UMM website.

The Dataset-PAT consists of 11,970 WSIs from 10 hospitals, ACL and the cancer genome atlas (TCGA-FFPE, https://portal.gdc.cancer.gov/), which were used for extensive patient-level prediction. The WSIs from 9 of 12 centers and 213 WSIs from XH were included to Dataset-HAC for human-AI competition after checking their labels carefully. Because XH was the biggest data source, the WSIs from XH were distributed independently and exclusively in Dataset-PATT, Dataset-PT and Dataset-HAC.

**Digitization and annotation of pathological slides**

In the 10 hospitals and ACL, the technicians randomly selected slides from archive library. The slides from 2010-2019 were scanned with a KF-PRO-005 scanner (KFBIO company, Ningbo City, China) at 20X magnification. The number of selected patients collected on the same day was limited to less than 50 to make sure the selected WSIs for this study were not unduly influenced by samples collected on any one single day.

All diagnosis of images from TCGA, NCT-UMM were available online, and their labels were used directly. The WSIs from the 10 hospitals and ACL in Dataset-PT were independently reviewed by two senior and seasoned pathologists. When their diagnoses were consistent, the WSI were then included. Dataset-HAC was used for human-AI competition, and the review criteria were more rigorous. The WSI label in Dataset-HAC were more strictly checked by three senior highly experienced pathologists who independently reviewed the pathological images without knowing the previous clinical diagnosis. If a consensus was reached, the WSI were included; otherwise, two other independent pathologists would join the review. After a discussion among the five pathologists, the WSI was included for the Human-AI contest only if they reached an agreement.

**Annotation of patches in Dataset-PATT**

The presented approach is based on the patch-level prediction. There is high phenotypic diversity within tumor and among tumors, the representation of cancer tissue in patches seriously affects the training. Therefore, the patches in Dataset-PATT were carefully selected to include all common tumor histological subtypes, ensuring the selected patches were widely representative for practical diagnosis.

The technician randomly selected 842 slides from pathological archive library of Xiangya hospital and then scanned them using a KF-PRO-005 scanner (KFBIO company, Ningbo City, China) at 20X magnification. Because the shape of the CRC tissue is less diverse than that of non-cancerous tissue, more cancer positive WSIs (n=614) and less cancer negative WSIs (n=228) were selected. For the 614 positive WSIs, the number of positive WSIs of various CRC subtypes were basically consistent with the subtype morbidity in the population.

Two pathologists used image browser software provided by KFRIO company (Ningbo City, China) from one WSI to export some non-overlapping regions of interest (ROI) according to the size of WSI. In order to maintain the diversity of cancer cell distribution, the 4-10 positive ROIs are extracted from each positive WSI. In order to ensure that the number of positive ROIs and negative ROIs is balanced, the 10-25 ROIs are extracted from each negative WSI. One ROI has a size of about $1024\times768$ pixels, and was split into about 6 non-overlapping patches with $300\times300$ pixels in order to be adaptable to meet the input size of most neural networks. The two pathologists then manually reviewed the patches, each of which was weakly labeled with either cancer or cancer-free. When two pathologists reached a consensus on the annotation of patches, which were kept in the Dataset-PATT.

In total, 62,919 patches were obtained. The 30,056 labeled tumor patches from 614 patients and 32,863 normal patches from 228 healthy subjects were included in Dataset-PATT, that is, an average of 49 patches per cancerous WSI and 144 patches per healthy WSI were included. Meanwhile, the number of patches containing various proportion of cancer cells were approximately equal.

**Patch-level SSL and SL models**

The Dataset-PATT was randomly divided into training set and testing set according to the proportions showed in Table 3, and the patches from the same subject would not be in different sets, to ensure independence of the different data sets. Meanwhile, the patches from 70% of the subjects were used as the training set, while the remaining 30% subjects were used as the testing set.

Five patch-level models (two SSL, three SL) were trained using labels of different portions of these patches (Table 3). In the training of Model-5%-SSL and Model-10%-SSL, we used SSL and kept labels for small proportions (i.e., 5% and 10%) of the total patches (62,919) and masked label information for the remaining patches (65% and 60%). In the training of Model-5%-SL, Model-10%-SL and Model-70%-SL, we used SL with 5%, 10%, 70% of the total 62,919 patches. Because training a deep learning model is time-consuming, for illustration, we repeated the process 8 times and calculated 0.75 confidence interval. The Model-70%-SL was trained on the same number of labeled patches with SL as in our previous study [2]. The Dataset-PAT was used as an independent test.

**Algorithm pipeline**

Because WSI is very large (>50,000 pixels), the patches in a WSI were firstly extracted, and the patch-level models were trained to derive cancerous probability at patch-level. Finally, all the patch-level results on a WSI were combined to infer the cancerous probability of the WSI/patient. The flow chart is shown in Figure 1.

**Patch-level SSL and SL**

The patch-level models included SL and SSL versions. For SL, the patches from the WSIs were input to the CNN. Our previous work tested some known CNNs, such as VGG16, ResNET V1 and V2, Inception V1-V4, and found that Inception V3 [1] achieved most consistent results on the CRC datasets [2]. Therefore, we used Inception V3 as the baseline model of SL. The patch size we labeled was 300×300, so we used the bilinear interpolation method to scale the patch size to 299×299, which is the default input size of Inception V3. The top output layer was removed, and the output category was modified to two (cancer or non-cancer).

The SSL version was implemented based on the mean teacher method [3], where two Inception V3 were trained, one as student and the other as teacher, which is one of SSL method (Supplementary Figure 1). The student network uses SL and requires inputted patches, which include a small number of patches with labels and large number of unlabeled patches. For the labeled patches, the cross-entropy of the predicted and real label was calculated as the classification cost. For unlabeled patches, teacher network provided the pseudo labels, and the mean square of the predicted labels and pseudo labels was calculated as consistency cost. The weighted sum of consistency cost and classification cost, as the total cost, was used for the student network training. In this study, the two networks were performed on the same architecture with SL, i.e., Inception V3.

**Network training at patch level**

The Inception V3 was initialized with a pre-trained model on ImageNet database, and then trained on the pathological images. During training, the weights in all layers of inception V3 were updated. We used the same preprocessing in protocols we used earlier [2]. All background patches without any cell tissue were removed. After data augmentation (image zoom, flip, color change), the grayscale of each pixel was normalized to [-1,1].

For each model, we adopted a general strategy where the one-tenth of the labeled training set was taken out as the validation set for hyperparameters selection. The optimal hyper-parameters with highest accuracy in the validation set were selected for training the models. The parameters were listed in Supplementary Table 2.

In the SSL, because of the imbalance between the labeled and unlabeled data, we maintained the same proportion of labeled and unlabeled patches in each mini-batch of 128 patches. The optimizer was Adam. The training period was 500 epochs, and each epoch included 100 steps. If the accuracy on validation set can't be improved for 80 consecutive epochs, the early stopping [22] was applied. In order to prevent the training from ending prematurely, 50 epochs for pre-training were executed before the early stopping. L2 decay was used and the decay coefficient was set to 0.0001. The teacher network was initialized with the student network. The student network would update the weights in each step, but the teacher network used exponential moving average to update the weights after one epoch ended. The smoothing coefficient was set to 0.95.

In SL, the learning rate was 0.001, and the exponentially decay was used with the decay rate 0.99. The number of epochs was 500, the steps per epoch was 100. The early stopping with patience 50 was also applied. The coefficient of L2 decay was 0.0001, and the batch size was 64.

**Clustered-based WSI inference**

Because the accuracy of patch-level models cannot be 100%, there were serious false positives in WSI predictions if any patch in the WSI was identified as positive (cancer) and used as a criterion for predicting the WSI cancerous status. Intuitively, because the tissues in WSI were continuous, the area with cancer should be distributed continuously and included several continuous positive patches. This intuition had been used to effectively control the false-positive of functional magnetic resonance images [4]. We designed a simple clustering-based inference method. If some continuous patches were identified as having cancer by patch-level model, the cancer may indeed exist on WSI. The cluster size of four patches was expected to best control the false-positive rate as shown in our early study [2], that is, the condition of continuously identifying 4 patches with cancer on WSI was used as the basis for determining the existence of cancer in WSI. For statistical analysis on patient-level prediction, please refer to Supplementary B.

**Patient-level diagnosis**

Clinically, multiple WSIs may be obtained for one patient. The inference on patient level was based on positive sensitivity, that is, if all WSIs from the same patient were identified as negative (no cancer), then the patient was negative, otherwise the patient was positive.

The source code and data can be found: www.github.com/csu-bme/pathology_SSL. The code was implemented in Python(version 3.6.9) [5] and Tensorflow (version 1.15.0) [6].

**B.** **Statistical analysis on patient-level prediction**

The patient-level diagnosis was based on two strategies: cluster-based WSI inference and positive sensitivity for patient inference. The main purpose of cluster-based WSI inference was to control the false positive rate (FPR) on each WSI, because the cancerous probability on each patch was not absolutely accurate, and multiple tests of many patches greatly increased false positives on one WSI.

Assuming the patch-level sensitivity and specificity were $\theta$ and $\gamma$, there were k consecutive positive patches on one WSI, i.e. the positive cluster size was $k$. At the same time, we also assumed that these patches were mutually independent. Theoretically, the probability of correctly identifying the WSI with one positive cluster was $\theta^{k}$, and the probability of falsely identifying one WSI of non-cancer was ${(1-\gamma)}^{k}$. Assuming $k=3$, $\gamma$=0.95, we had a patch-level FPR=0.05, while the FPR≈0.0001 on the WSI.

In practice, adjacent patches on a WSI were highly correlated. Consequently, the above theoretical derivation was not precise enough. However, the experiments proved that the false positive control of one WSI can achieve high predictive power with a cluster of four positive contiguous patches [2]. Therefore, we used the clustering of 4 patches as the condition for positive WSI inference. Finally, as long as the patient had a positive WSI, he or she was diagnosed with CRC (positive sensitivity).

**C.** **Patient-level Comparison of Model-10%-SSL, Model-10%-SL, Model-70%-SL**

Patient-level comparisons were performed on Model-10%-SSL, Model-10%-SL, Model-70%-SL. The data source name, as well as the sensitivity, specificity, accuracy, and AUC values were given in Supplementary Table 3. The three models were trained in SSL or SL at patch-level, but the same strategies of cluster-based WSI inference and positive sensitivity for patient inference were used at the patient level.

**D.	Methodology of extended lung and lymph node**

Two public datasets were used for the extended evaluation of SSL. The Dataset-Lung was from the LC25000 [7], which consisted of 15,000 lung images (patches) including adenocarcinoma, squamous cell carcinoma, and benign tissue, and the number of each class was 5,000 patches. The 20% images were used for testing, while the remaining 80% for training. A small number of labels (5%, 20% of 15,000 patches) together with large number of unlabeled patches (75%, 60%, the labels were ignored) were used for SSL, while the 5%, 20% and 80% labeled images were used for SL (Supplementary Table 6).

The Dataset-Pcam was from PatchCamelyon dataset [8] including up to 300,000 patches of lymph node tissues, which had been split into training, validation and testing sets. Meanwhile, the number of patches in training set was 262,144 patches, and the 1% and 5% of the patches was randomly extracted to simulate a small number of labeled data, while the remaining patches to simulate massive unlabeled data (the labels were ignored). The 36,728 patches in testing set were used for testing (Supplementary Table 7), but 32,768 patches of validation set were not used.

In order to evaluate the statistics, the experiments were randomly repeatedly extracted 8 times. Like CRC experiments, the base SL model was also Inception V3, and the mean teacher method was used for SSL. The 10% patches of labeled training set were randomly selected for the validation set, which was used for the hyperparameter selection. This selection starts from the hyperparameters of the CRC models, and tries the parameters nearby. The parameters are listed in Supplementary Table 8 and 9.

The processing pipeline of the images from the lung and lymph node was like the CRC. Meanwhile, the patches are scaled to $299\times299$ based on the image interpolation. For SL of lung, the batch size was 64, the number of epochs was 500, the steps of each epoch was 100. The initial learning rate was 0.001, and the exponential decay was used with the decay rate was 0.99. The loss is the cross entropy with the L2 norm constraint, whose the coefficient the L2 decay was equal to 0.0001. The early stopping was also used, where the patience was 50 epochs. For SSL of lung, the batch size was 32, the number of epochs, the steps, loss were the same with SL, but the learning rate was 0.0001, and remained the same. After the pre-training of 150 epochs, the early stopping was also used with the patience of 100 epochs. The smoothing coefficient of exponential moving average of teacher network and student network is set to 0.9.

Because the number of training patches in Dataset-Pcam was very large and the experiment time was very long, we continued to use the hyperparameters in lung experiments and tried to optimize them. For SL, the batch size, epochs, initial learning rate, decay rate, weights of L2 decay were the same with SL of lung, but the steps were changed to 300. We found the AUC and accuracy of Pcam-100%-SL can be compared to the benchmark provided by [8], so the hyperparameters were applicable. For SSL, the steps were 200. After the training of 80 epochs, the early stopping with patience 100 was used. The remaining hyperparameter were the same with SSL for lung models.

**Supplementary Table 1. Allocation (number) of CRC WSIs from 13 data centers**

| **Dataset** | **PATT** | **PAT** | **PT** | **HAC** |
| --- | --- | --- | --- | --- |
| XH | 842 | 0 | 10003 | 213 |
| NCT-UMM | 0 | 86 | 0 | 0 |
| TXH | 0 | 0 | 135 | 135 |
| PCH | 0 | 0 | 96 | 96 |
| HPH | 0 | 0 | 99 | 99 |
| FUS | 0 | 0 | 198 | 198 |
| GPH | 0 | 0 | 185 | 185 |
| SWH | 0 | 0 | 199 | 199 |
| AMU | 0 | 0 | 205 | 205 |
| SYU | 0 | 0 | 97 | 97 |
| ACL | 0 | 0 | 207 | 207 |
| CGH | 0 | 0 | 100 | 0 |
| TCGA | 0 | 0 | 446 | 0 |
| **Total** | 842 | 86 | 11,970 | 1,634 |

**Supplementary Table 2. Hyper-parameters used in SSL and SL of CRC**

| SSL |  |
| --- | --- |
| Hyper-parameters | **Value** |
| Learning rate | 0.0001 |
| Optimizer | Adam |
| Epochs | 500 |
| Step per epoch | 100 |
| Batch size | 128 |
| L2 decay | 0.0001 |
| Pre trained epochs | 50 |
| Early stopping | True |
| Patience | 80 |
| smoothing coefficient | 0.95 |
| SL | |
| Hyper-parameters | **Value** |
| Learning rate | 0.001 |
| decay rate | 0.99 |
| Optimizer | Adam |
| Epochs | 500 |
| Batch size | 64 |
| Step per epoch | 100 |
| L2 decay | 0.0001 |
| Early stopping | True |
| Patience | 50 |

**Supplementary Table 3. Patient-level Comparison of Model-10%-SSL, Model-10%-SL, Model-70%-SL**

| MOLEs | | Data source | Sensitivity | | Specificity | | Accuracy | | AUC | | AUC-DIFF | |
| --- | --- | --- | --- | --- | --- | --- | --- | --- | --- | --- | --- | --- |
| Model-10%-SSL | | XH-Dataset-PT | 0.989 | | 0.979 | | 0.986 | | 0.984 | | -0.008 | |
|  | | XH-Dataset-HAC | 0.990 | | 0.949 | | 0.969 | | 0.984 | | -0.006 | |
|  | | PCH | 0.960 | | 0.848 | | 0.906 | | 0.954 | | -0.015 | |
|  | | TXH | 1.000 | | 0.958 | | 0.979 | | 0.979 | | -0.010 | |
|  | | HPH | 1.000 | | 0.980 | | 0.990 | | 0.990 | | **0.010**** | |
|  | | FUS | 0.990 | | 0.908 | | 0.950 | | 0.974 | | -0.016 | |
|  | | GPH | 0.980 | | 0.929 | | 0.956 | | 0.966 | | **-0.022*** | |
|  | | SWH | 0.970 | | 0.930 | | 0.950 | | 0.974 | | -0.016 | |
|  | | SYU-CGH^a^ | 0.978 | | 0.906 | | 0.939 | | 0.966 | | **0.009**** | |
|  | | AMU | 0.960 | | 0.961 | | 0.961 | | 0.984 | | **0.014**** | |
|  | | ACL | 0.940 | | 0.935 | | 0.937 | | 0.946 | | **-0.040*** | |
|  | | TCGA-FFPE | 0.972 | | 1.000 | | 0.973 | | 0.986 | | -0.004 | |
|  | | Average | 0.977 | | 0.940 | | 0.958 | | 0.974  $P \mathrm{value}=0.117$ | | -0.006 | |
| MODEL-10%-SL | | XH-Dataset-PT | 0.993 | | 0.936 | | 0.974 | | 0.964 | | **-0.028*** | |
|  | | XH-Dataset-HAC | 0.776 | | 0.876 | | 0.826 | | 0.826 | | **-0.164*** | |
|  | | PCH | 0.918 | | 0.739 | | 0.832 | | 0.829 | | **-0.140*** | |
|  | | TXH | 1.000 | | 0.833 | | 0.917 | | 0.917 | | **-0.073*** | |
|  | | HPH | 1.000 | | 0.265 | | 0.633 | | 0.633 | | **-0.347*** | |
|  | | FUS | 1.000 | | 0.214 | | 0.611 | | 0.633 | | **-0.357*** | |
|  | | GPH | 1.000 | | 0.718 | | 0.869 | | 0.877 | | **-0.111*** | |
|  | | SWH | 0.980 | | 0.680 | | 0.829 | | 0.869 | | **-0.111*** | |
|  | | SYU-CGH | 1.000 | | 0.349 | | 0.650 | | 0.717 | | **-0.240*** | |
|  | | AMU | 1.000 | | 0.520 | | 0.754 | | 0.782 | | **-0.188*** | |
|  | | ACL | 0.840 | | 0.880 | | 0.860 | | 0.859 | | **-0.127*** | |
|  | | TCGA-FFPE | 0.843 | | 1.000 | | 0.845 | | 0.922 | | **-0.068*** | |
|  | | Average | 0.946 | | 0.668 | | 0.800 | | 0.819,  $P \mathrm{value}=0.002$ | | **-0.161*** | |
|  | MODEL-70%-SL | XH-Dataset-PT | | 0.988 | | 0.995 | | 0.990 | | 0.992 | | - |
|  |  | XH-Dataset-HAC | | 0.980 | | 1.000 | | 0.990 | | 0.990 | | - |
|  |  | PCH | | 0.960 | | 0.978 | | 0.969 | | 0.969 | | - |
|  |  | TXH | | 1.000 | | 0.979 | | 0.990 | | 0.990 | | - |
|  |  | HPH | | 0.980 | | 0.980 | | 0.980 | | 0.980 | | - |
|  |  | FUS | | 1.000 | | 0.980 | | 0.990 | | 0.990 | | - |
|  |  | GPH | | 1.000 | | 0.977 | | 0.989 | | 0.988 | | - |
|  |  | SWH | | 0.990 | | 0.970 | | 0.980 | | 0.980 | | - |
|  |  | SYU-CGH | | 0.989 | | 0.925 | | 0.954 | | 0.957 | | - |
|  |  | AMU | | 0.970 | | 0.971 | | 0.970 | | 0.970 | | - |
|  |  | ACL | | 1.000 | | 0.972 | | 0.986 | | 0.986 | | - |
|  |  | TCGA-FFPE | | 0.980 | | 1.000 | | 0.980 | | 0.990 | | - |
|  |  | Average | | 0.982 | | 0.977 | | 0.979 | | 0.980 | | - |

AUC-DIFF= Model-10%-SSL/AUC – Model-70%-SL/AUC

= Model-10%-SL/AUC – Model-70%-SL/AUC

*: AUC-DIFF<-0.016; **: AUC-DIFF is positive value

a: Because the positive and negative samples a of SYU and CGH were unbalanced, their metrics were calculated together.

The $P\mathrm{value}$ was the Wilcoxon signed rank test result between Model-70%-SL/AUC and Model-10%-SSL/AUC or Model-10%-SL/AUC.

**Supplementary Table 4. Pathologist info full spelling here**

| **Pathologist ID** | **Years in Clinic** | **Job Title** |
| --- | --- | --- |
| A | 1 | Resident physician |
| B | 3 | Resident physician |
| C | 5 | Physician-in-charge |
| D | 7 | Physician-in-charge |
| E | 12 | Physician-in-charge |
| F | 18 | Associate chief physician |

**Supplementary Table 5. Overall performance of two models and pathologists in Human-AI competition**

|  | Model-10%-SSL | Model-70%-SL | Pathologists | | | | | | |
| --- | --- | --- | --- | --- | --- | --- | --- | --- | --- |
|  |  |  | A | B | C | D | E | F | Average |
| Sensitivity | 0.977 | 0.987 | 0.914 | 0.945 | 0.988 | 0.992 | 0.981 | 0.953 | 0.962 |
| Specificity | 0.930 | 0.980 | 0.981 | 0.975 | 0.965 | 0.982 | 0.986 | 0.960 | 0.975 |
| Accuracy | 0.954 | 0.983 | 0.955 | 0.967 | 0.975 | 0.986 | 0.984 | 0.956 | 0.971 |
| AUC | 0.972 | 0.984 | 0.951 | 0.960 | 0.977 | 0.987 | 0.983 | 0.956 | 0.969 |
| AUC RANK | 5 | 2 | 9 | 7 | 4 | 1 | 3 | 8 | 6 |

**Supplementary** Table 6. Training and testing set for lung models

| Model | Class | Dataset-Lung | | |
| --- | --- | --- | --- | --- |
|  |  | Training set | | Testing set |
|  |  | Labeled | Unlabeled |  |
| Lung-5%-SSL | adenocarcinoma | 250 | 3750 | 1,000 |
|  | squamous cell carcinoma | 250 | 3750 | 1,000 |
|  | benign | 250 | 3750 | 1,000 |
|  | Total | 750/5% | 11,250/75% | 3,000/20% |
| Lung-20%-SSL | adenocarcinoma | 1,000 | 3,000 | 1,000 |
|  | squamous cell carcinoma | 1,000 | 3,000 | 1,000 |
|  | benign | 1,000 | 3,000 | 1,000 |
|  | Total | 3,000/20% | 9,000/60% | 3,000/20% |
| Lung-5%-SL | adenocarcinoma | 250 | - | 1,000 |
|  | squamous cell carcinoma | 250 | - | 1,000 |
|  | benign | 250 | - | 1,000 |
|  | Total | 750/5% | - | 3,000/20% |
| Lung-20%-SL | adenocarcinoma | 1,000 | - | 1,000 |
|  | squamous cell carcinoma | 1,000 | - | 1,000 |
|  | benign | 1,000 | - | 1,000 |
|  | Total | 3,000/20% | - | 3,000/20% |
| Lung-80%-SL | adenocarcinoma | 4,000 | - | 1,000 |
|  | squamous cell carcinoma | 4,000 | - | 1,000 |
|  | benign | 4,000 | - | 1,000 |
|  | Total | 12,000/80% | - | 3,000/20% |

**Supplementary** Table 7. Training and testing set for lymph node models

| Model | Class | Dataset-Pcam | | |
| --- | --- | --- | --- | --- |
|  |  | Training set | | Testing set |
|  |  | Labeled | Unlabeled |  |
| Pcam-1%-SSL | Tumor | 1,311 | 129,761 | 16,384 |
|  | Non-tumor | 1,311 | 129,761 | 16,384 |
|  | Total | 2,622/1%^a^ | 259,522/99%^b^ | 32,768 |
| Pcam-5%-SSL | Tumor | 6,554 | 124,518 | 16,384 |
|  | Non-tumor | 6,554 | 124,518 | 16,384 |
|  | Total | 13,108/5%^c^ | 249,036/95%^d^ | 32,768 |
| Pcam-1%-SL | Tumor | 1,311 | - | 16,384 |
|  | Non-tumor | 1,311 | - | 16,384 |
|  | Total | 2,622/1% | - | 32,768 |
| Pcam-5%-SL | Tumor | 6,554 | - | 16,384 |
|  | Non-tumor | 6,554 | - | 16,384 |
|  | Total | 13,108/5% | - | 32,768 |
| Pcam-100%-SL | Tumor | 131,072 | - | 16,384 |
|  | Non-tumor | 131,072 | - | 16,384 |
|  | Total | 262,144/100%^e^ |  | 32,768 |

a-e: 1%, 99%, 5%, 95%, 100% of training set.

**Supplementary Table 8. Hyper-parameters used in SSL and SL of lung**

| SSL |  |
| --- | --- |
| Hyper-parameters | **Value** |
| Learning rate | 0.0001 |
| Optimizer | Adam |
| Epochs | 500 |
| Steps per epoch | 100 |
| Batch size | 32 |
| L2 decay | 0.0001 |
| Pre trained epochs | 150 |
| Early stopping | True |
| Patience | 100 |
| smoothing coefficient | 0.9 |
| SL | |
| Hyper-parameters | **Value** |
| Learning rate | 0.001 |
| decay rate | 0.99 |
| Optimizer | Adam |
| Epochs | 500 |
| Steps per epoch | 100 |
| Batch size | 64 |
| L2 decay | 0.0001 |
| Early stopping | True |
| Patience | 50 |

**Supplementary Table 9. Hyper-parameters used in SSL and SL of** lymph node

| SSL |  |
| --- | --- |
| Hyper-parameters | **Value** |
| Learning rate | 0.0001 |
| Optimizer | Adam |
| Epochs | 500 |
| Steps per epoch | 200 |
| Batch size | 32 |
| L2 decay | 0.0001 |
| Pre trained epochs | 80 |
| Early stopping | True |
| Patience | 100 |
| smoothing coefficient | 0.9 |
| SL | |
| Hyper-parameters | **Value** |
| Learning rate | 0.001 |
| decay rate | 0.99 |
| Optimizer | Adam |
| Epochs | 500 |
| Steps per epoch | 300 |
| Batch size | 64 |
| L2 decay | 0.0001 |
| Early stopping | True |
| Patience | 50 |

**Supplementary Table 10. List of AUCs of AI applied in CRC and other cancer types**

| Study | Patch-level test data | | Independent patch-level test data | | Slide-level test data | | Independent slide-level test data | | |
| --- | --- | --- | --- | --- | --- | --- | --- | --- | --- |
|  | Number (#) of patches | AUC | # of patches | AUC | # of slides | AUC | # of datasets | # of slides | AUC |
| **Colorectal cancer** | | | | | | | | | |
| Haj‑Hassan et al.^[9]^ | NA | Unsegmented~0.7923  Segmented~0.9917 | NA | NA | NA | NA | NA | NA | NA |
| Xu et al.^[10]^ | 717 | 0.969-0.980^a^ | NA | NA | NA | NA | NA | NA | NA |
| Sari et al.^[11]^ | 1,592 | 0.994 | NA | NA | NA | NA | NA | NA | NA |
| Kainz et al.^[12]^ | 60 | 0.983^a^ | 20^a^ | 0.950^a^ | NA | NA | NA | NA | NA |
| Kather et al.^[13]^ | 100,000 | 0.987 | 7,180 | 0.943 | NA | NA | NA | NA | NA |
| Ponzio et al.^[14]^ | 4500 | 0.9037-0.9682 | NA | NA | NA | NA | NA | NA | NA |
| Shaw et al.^[15], c^ | 7,180 | 0.9377 | NA | NA | NA | NA | NA | NA | NA |
| **Model-10%-SSL** | About 20,000 | 0.99 | 100,000 | 0.971 | 10,216^d^ | 0.984 | 11 | 1,967 | 0.946-0.99 |
| **Model-70%-SL** | About 20,000 | 0.994 | 100,000 | 0.98 | 10,216 | 0.989-0.992 | 11 | 1,967 | 0.96-0.99 |
| **Other cancers** | | | | | | | | | |
| Coudray et al.^[16]^ /lung cancer | NA | NA | NA | NA | 244 | 0.990-0.993 | 3 | 340 | LUAD~0.833-0.913  LUSC~0.861-0.941 |
| Cruz-Roa et al.^[17]^ / ductal carcinoma | 50,963 | 0.842^b^ | NA | NA | NA | NA | NA | NA | NA |
| Araujo et al.^[18]^ /breast cancer | 240 | 0.829^a^ | 192 | 0.693^a^ | 20 | 0.900^a^ | 1 | 16 | 0.750^a^ |
| Motlagh et al.^[19]^/breast cancer | 2,147 | 0.999 | NA | NA | NA | NA | NA | NA | NA |
| Campanella et al.^[20], e^ | NA | NA | NA | NA | 12,132 | 0.986-0.991 | 1 | 12,727 | 0.986-0.991 |
| Campanella et al.^[20]^ | NA | NA | NA | NA | 6,252 | 0.986-0.988 | 1 | 3,710 | 0.986-0.988 |
| Campanella et al.^[20]^ | NA | NA | NA | NA | 8,670 | 0.965-0.966 | 1 | 1,224 | 0.965-0.966 |

Note: a: accuracy; b: balanced accuracy; c: semi-supervised learning, accuracy on training sets with 20% labeled data; d: XH-Dataset-PT and XH-Dataset-HAC.

e: AUC for prostate cancer, basal cell carcinoma and breast cancer metastases

real label

consistency cost

pseudo label

Teacher network

Student network

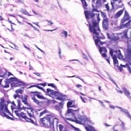

classification cost

cancer

non-cancer

cancer

non-cancer

predicted label

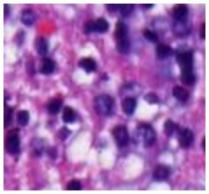

Labeled patches

Unlabeled patches

cancer

non-cancer

Supplementary Figure 1. The flow chart of mean teacher.

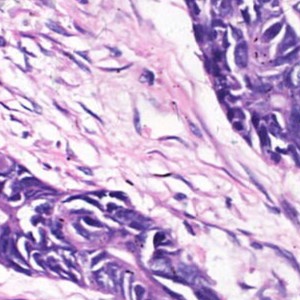

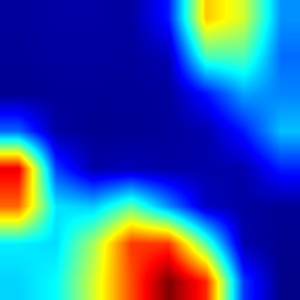

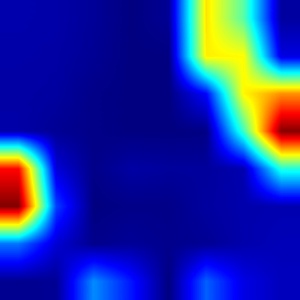

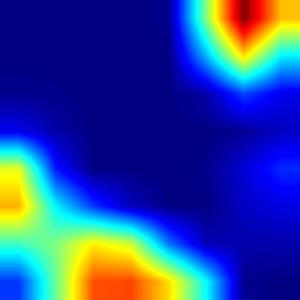

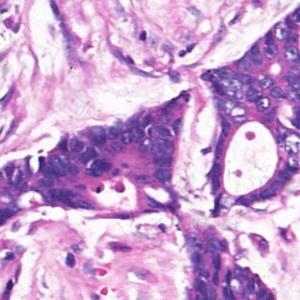

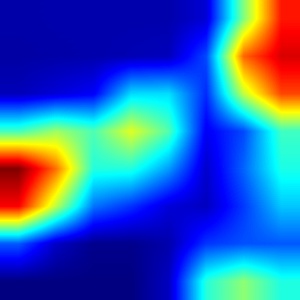

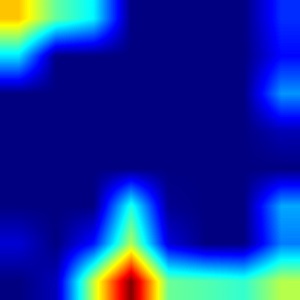

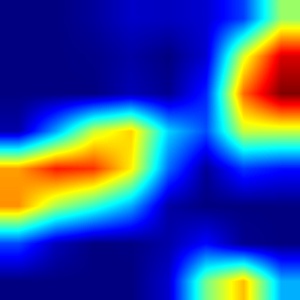

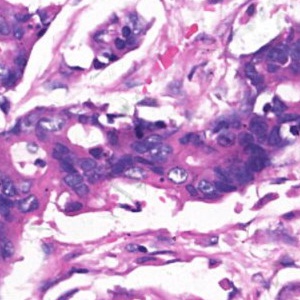

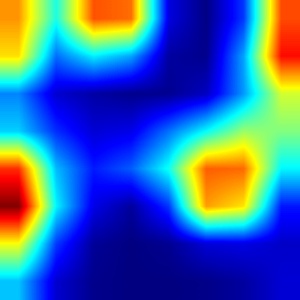

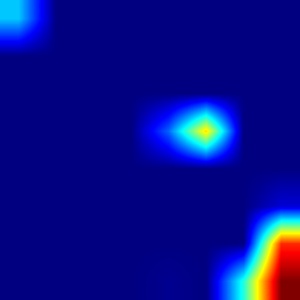

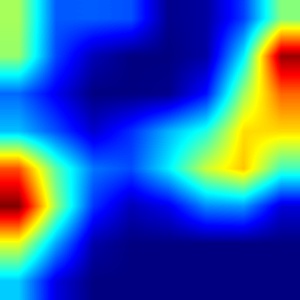

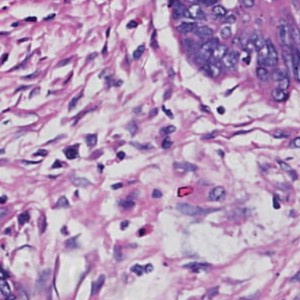

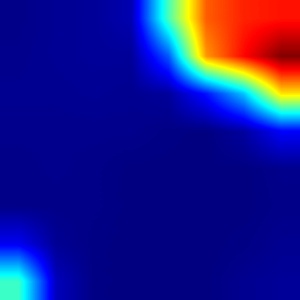

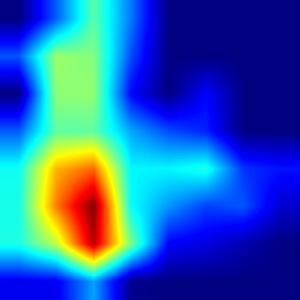

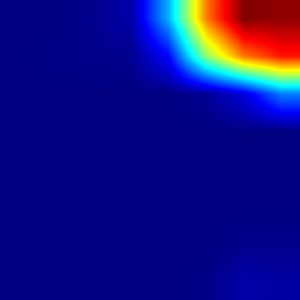

Original image	Model-10%-SSL	Model-10%-SL	Model-70%-SL

Supplementary Figure 2. The heatmaps on cancerous pixels probability in patches. Some representative patches were selected and the pixels that have an important contribution to identifying cancers are shown with colors. The more important the pixel, the warmer the color. It can be seen that the activated regions of Model-10%-SSL and Model-70%-SL are very similar to each other, and the cancerous pixels are highly overlapped with most of the tumor cell areas. However, the heatmaps of Model-10%-SL are deviated from that of Model-10%-SSL and Model-70%-SL, both the size and location of cancerous regions in some patches (row 2,3,4).

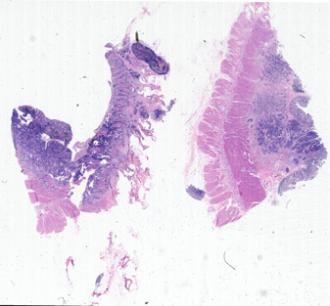

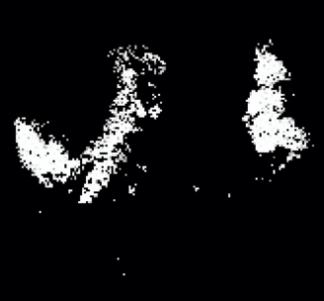

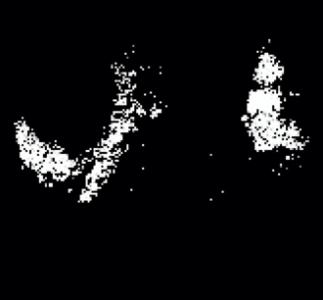

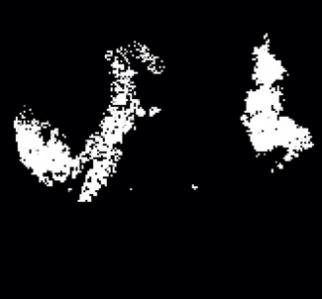

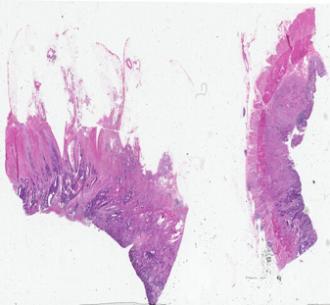

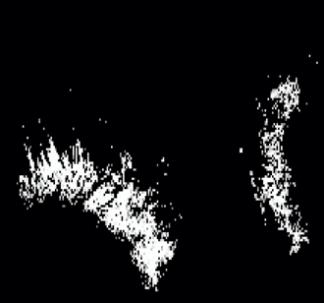

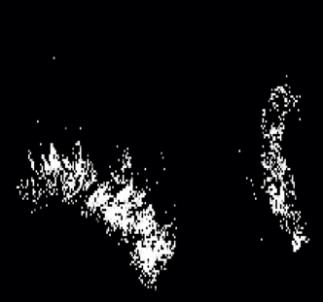

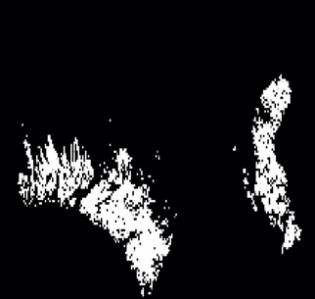

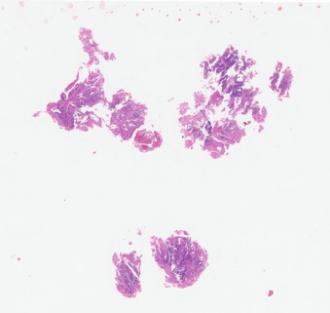

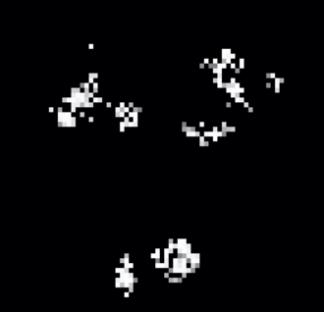

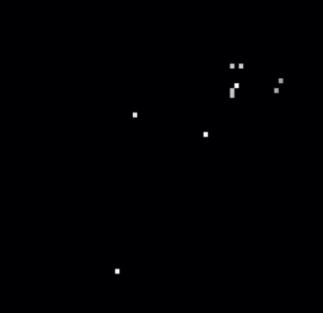

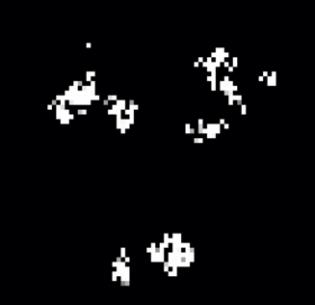

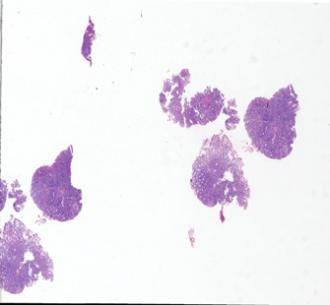

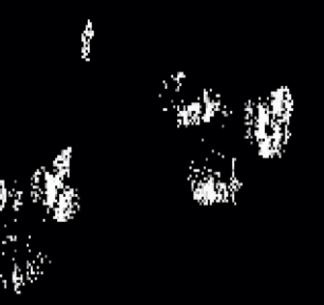

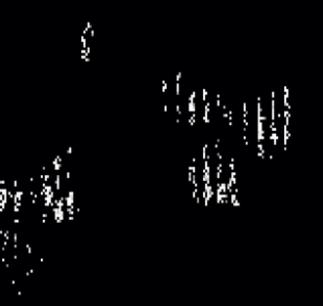

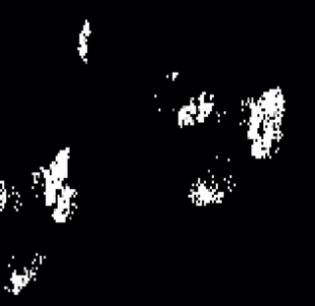

Whole slide image	Model-10%-SSL Model-10%-SL Model-70%-SL

Supplementary Figure 3: The cancerous regions on the WSI predicted by Model-10%-SSL, Model-10%-SL, and Model-70%-SL are shown in white. The white regions of Model-10%-SSL and Model-70%-SL are similar to each other and highly overlapped with the most area given by pathologists. However, the results of Model-10%-SL are deviated from that of Model-10%-SSL and Model-70%-SL, especially for the WSIs from the colonoscopy (row 3 and 4).

**Reference**

[1] Christian Szegedy, Vincent Vanhoucke, Sergey Ioffe, Jonathon Shlens, Rethinking the Inception Architecture for Computer Vision, arXiv preprint arXiv: 1512.00567v3.

[2]. Wang Kuan-Song, Yu Gang, Xu Chao, et al, Accurate Diagnosis of Colorectal Cancer Based on Histopathology Images Using Artificial Intelligence. BMC Medicine, 2021, 19(76):1-12.

[3]. Antti Tarvainen, Harri Valpola, Mean teachers are better role models: Weight-averaged consistency targets improve semi-supervised deep learning results, arXiv preprint arXiv:1703.01780v6.

[4].	Heller R, Stanley D, Yekutieli D, Rubin N, Benjamini Y. Cluster-based analysis of FMRI data. Neuroimage 2006, 33:599-608.

[5] Python Software Foundation. Python. Version 3.6.9[software]. 2019 June 18 [cited 2020 Nov 23]. Available from: https://www.python.org.

[6] Google Inc. Tensorflow. Version 1.15.0[software]. 2019 Oct 17[cited 2020 Nov 23]. Available from: https://pypi.org/project/tensorflow.

[7] Borkowski AA, Bui MM, Thomas LB, Wilson CP, DeLand LA, Mastorides SM. Lung and Colon Cancer Histopathological Image Dataset (LC25000). arXiv:1912.12142v1 [eess.IV], 2019

[8] PatchCamelyon dataset, <https://github.com/basveeling/pcam>

[15] Shayne Shaw, Maciej Pajak, Aneta Lisowska, Sotirios A. Tsaftaris, Alison Q. ONel, Teacher-student chain for efficient semi-supervised histology image classification, arXiv preprint arXiv:2003.08797v2, 2020

[22] Prechelt L. Neural Networks: Tricks of the Trade. Springer Berlin Heidelberg, 1998.
